# Unpacking the eagle collision risk model: practical guidance for wind energy

**DOI:** 10.1101/2020.07.02.182907

**Authors:** Michael J. Evans, Misti Sporer, Wally Erickson, Joy Page

## Abstract

Climate change is one of the greatest threats facing biodiversity, and solutions to reduce carbon emissions are needed to conserve species. Renewable energies are a prominent means to achieve this goal, but the potential for direct harm to wildlife has raised concerns as these technologies proliferate. To protect biodiversity, approaches that facilitate renewable energy development while protecting species are needed. In the United States wind energy developers must obtain a permit for any Bald or Golden eagles that might be killed at a facility. The U.S. Fish & Wildlife Service estimates fatalities using a Bayesian modeling framework, which combines pre-construction eagle surveys with prior information. The ways in which prior information is incorporated and how pre-construction monitoring affects model outcomes can be unclear to regulated entities and other stakeholders, creating uncertainty in the permitting process and retarding both the build-out of renewable energy and conservation measures for eagles. We conducted a simulation study quantifying the differences in predicted eagle fatalities obtained by incorporating prior information and using only site-specific survey data across a range of scenarios, evaluating the impact of survey effort on the magnitude of this effect. We identified predictable relationships between survey effort, eagle activity, facility size and discrepancies between estimates. We also translated these patterns into real-world financial costs, illustrating the interaction between pre-construction surveys, fatality estimates, and compensatory mitigation obligations in determining permit timing and expense.

## Introduction

Climate change is one of the primary human drivers of biodiversity loss and decline, and unless greenhouse gas emissions are drastically reduced millions of species will face extinction (Díaz et al. 2019), many in the coming decades (Trisos et al. 2020). A transition from fossil fuels to renewable energy sources, especially wind and solar, as a means to reduce carbon emissions (Rogelj et al. 2018) is therefore critical to conserve biodiversity. Renewable energy generation is growing rapidly both in the United States and across the globe (Carley et al. 2017). In 2018, U.S. renewable energy capacity (249 GW) surpassed 20% of total electric capacity, and 42.9% of all new U.S. capacity additions (Koebrich et al. 2018). However, the proliferation of renewables has raised concerns that these technologies may have direct, negative impacts on wildlife (Allison et al. 2019; Rehbein et al. 2020). In some cases, existing conservation policies place renewable energy development at odds with wildlife conservation, juxtaposing two endeavors that should be complimentary with regards to protecting biodiversity. To maximize the preservation and restoration of biodiversity, a synergistic approach to conservation policy is needed – one that both minimizes direct mortality and habitat loss while facilitating the critical long-term benefits of reduced carbon emissions.

Unfortunately, policies regulating renewable energy and wildlife conservation have often been developed independently (Köppel et al. 2014). For example, the Bald and Golden Eagle Protection Act (BGEPA) was passed before widespread development of utility-scale wind energy (*Bald and Golden Eagle Protection Act* 1940). The statute outlaws the intentional or unintentional ‘take’ (i.e. harm, harassment, killing) of Bald (*Haliaeetus leucocephalus*) or Golden eagles (*Aquila chrysaetos*) without a permit. Because wind energy can negatively affect eagles in a variety of ways (Johnson et al. 2016), this has placed wind energy development at odds with eagle conservation in the United States. Golden eagles are of particular concern; they may be declining in parts of their range (Millsap et al. 2013) and wind energy can create population sinks in certain regions (Katzner et al. 2017). As the goal of conservation should be to both reduce wildlife mortality and transition to renewable energy, policies aligning these goals are required.

To address this need, the U.S. Fish and Wildlife Service (hereafter ‘the Service’) developed guidance to harmonize wind energy facility development with eagle conservation through a permitting process (U.S. Fish and Wildlife Service 2013). The Service can issue permits under BGEPA authorizing eagle take for various approved activities (e.g., scientific collecting or research, religious purposes, falconry, etc.) including incidental take that occurs as a result of otherwise legal action. Wind energy operators are encouraged – but not required - to obtain an eagle take permit when building or operating a facility that might kill bald or golden eagles. To obtain a permit, operators are required to minimize and mitigate impacts to eagles to the extent practicable. At facilities that will be unable to avoid killing golden eagles, operators must offset these fatalities through mitigation. To date, this has often involved retrofitting high-risk power poles to prevent eagle electrocution on electric distribution lines, which is a leading anthropogenic cause of eagle mortality (U.S. Fish and Wildlife Service 2016). In principle these policies provide a means to develop wind energy capacity while minimizing impacts to a protected species. In practice, questions and concerns with the structure, execution, predictability, and timing of the Service’s permitting approach have led to fewer permits than possible.

At the core of this consternation is the Service’s use of a Bayesian framework to predict eagle fatalities at proposed wind energy facilities, known as the collision risk model. In this model, eagle survey data from existing wind facilities are incorporated with pre-construction survey data at a proposed site to estimate how many eagles are likely to be killed (New et al. 2015). The incorporation of this prior information is both an advantage of using a Bayesian framework (Ellison 2004) and a source of uncertainty and friction for operators, because it means a given facility’s predicted eagle fatalities may not be based on conditions at that site alone. This is problematic for operators because these predictions determine real mitigation costs for a facility (by influencing the estimate of eagle fatalities). The development of a wind site is a complex process with many stakeholders and overlapping requirements (e.g., local, state, federal regulations; environmental, financial, and engineering constraints; etc.). Unpredictable outcomes from known regulatory obligations contribute uncertainty to the costs and timelines associated with this process. If prior information is perceived to create uncertain, arbitrary outcomes for an operator they may avoid the eagle take permitting process altogether. Indeed, the wind industry has stated that eagle take permitting has unfairly slowed the development of wind energy (Speerschneider 2018). To date, less than 20 incidental take permits covering wind energy operation have been issued (personal comm.). If current policies unnecessarily slow wind development or incentivize developers to eschew the permitting process, then these policies are achieving suboptimal outcomes for both renewable energy and eagle conservation. A more certain process could accelerate wind energy and generate more eagle take permits, increasing conservation measures for eagles.

Here we illustrate and quantify the behavior of the collision risk model in order to help identify possible solutions for permitting that synergistically optimize eagle conservation while expediting renewable energy development. We focus our analyses on golden eagles, which are a species of conservation concern in the U.S. Our first objective was to demonstrate how eagle exposure priors interact with site-specific survey data to predict eagle fatalities. Second, we quantify how survey effort and wind energy facility characteristics change this relationship and affect fatality predictions. Finally, we evaluate and compare costs related to pre-construction eagle surveys and mitigation across a range of survey efforts and facility sizes. We used a simulated dataset containing a range of possible scenarios based on empirical data from wind farms (Bay et al. 2016). Our results clarify the behavior of the collision risk model and effects of survey effort on predicted fatalities, provide a quantitative framework for selecting minimum survey criteria, and may have utility as a means of identifying low-risk settings for expedited permitting.

## Methods

### Collision Risk Model

The eagle take permitting program under BGEPA has regulatory requirements for conducting the impact analysis that informs the required mitigation. The annual eagle fatalities at a proposed wind energy facility are estimated from three components:

1. *Eagle Exposure* (*min/hr*km^3^*): time eagles are estimated to fly within risk areas around turbines where collisions may occur.
2. *Collision Rate (fatalities/min)*: rate at which eagles flying within a turbine risk area are struck and killed.
3. *Expansion Factor* (*hr*km^3^*): scaling factor converting collision rates to annual fatalities based on the size and operating hours of a wind facility.

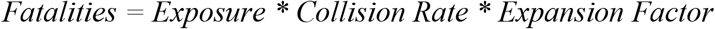

Within the Bayesian framework of the collision risk model, eagle exposure and collision rates are represented as probability distributions. Eagle collision rates are estimated entirely based on data from existing facilities. Eagle exposure is estimated from both survey data collected at the site of a proposed wind energy facility and a prior distribution of eagle exposure based on data collected from a sample of existing wind energy facilities. Survey data is combined with this prior distribution to create a posterior probability distribution of eagle exposure at a site (New et al. 2015). This posterior exposure distribution is multiplied by the global collision risk distribution and scaled by a site’s expansion factor to produce a distribution providing the probability that a given number of eagles will be killed annually, rather than a single point estimate (e.g., 80% chance that 5 or fewer eagles will be killed per year). The Service uses the 80^th^ percentile of this distribution as a facility’s annual fatality estimate to guard against underestimating eagle mortality (U.S. Fish and Wildlife Service 2013). In all analyses, we use the 80^th^ percentile as an estimate of eagle fatalities unless otherwise noted.

### Simulation data

To investigate the effects of prior information and survey effort on eagle fatality predictions, we created a simulation dataset representing a range of possible pre-construction survey values for survey area (km^3^), time (hr), and eagle observations (min). We then estimated annual eagle fatality for these scenarios using the collision risk model. To create the dataset we first generated a set of possible survey efforts. To obtain an eagle take permit, the Service requires 24 hours of survey effort over two years during which eagle minutes within at least one cylindrical survey plot are recorded. Plots are recommended to have an 800 m radius and be 200 m high. The minimum survey effort is therefore 0.402 hr*km^3^. We created a sample of potential survey efforts as the cartesian product of a set of number of survey plots {1, 2, 3, 4, 5} and a set of potential survey times {24, 48,…, 240 hr}, resulting in 50 levels of survey effort ranging from 0.402 - 482.4 hr*km^3^. Having created this set of survey efforts, we generated a set of potential eagle exposures (min) that would be recorded during surveys. Eagle exposure is a product of the intrinsic frequency with which eagles are active in the area (min/hr*km^3^, hereafter ‘eagle activity’) and the amount of survey effort (hr*km^3^). We generated a representative set of eagle activity rates {0.00, 0.05,…, 3.00 min/hr*km^3^}and created a final set of scenarios as the cartesian product of this set of rates and the set of survey efforts. Observed eagle exposure was then determined by multiplying the simulated eagle activity rate by the simulated survey effort for each scenario.

### Model behavior

To evaluate the effect of prior information on fatality estimates, we used the collision risk model to estimate fatalities incorporating the Service’s exposure priors as in New et al. (2015) and using only survey data for each simulated scenario. We refer to the difference in these estimates as “estimate discrepancy.” In all analyses, we used the collision risk model priors currently used by the Service. The prior distribution for eagle collision risk is ~ B (2.31, 396.69). The prior distribution of eagle exposure is ~ *Γ* (0.968, 0.552). A site-specific exposure distribution was defined as ~ *Γ* (eagle mins, survey effort). We generated the posterior distribution of predicted fatalities at a given site by multiplying 10,000 random draws from the relevant exposure distribution by 10,000 random draws from the collision risk prior distribution. We used the expansion factor (*ε*) to convert eagle fatality rates to predicted annual fatalities based on the size and operating time of a hypothetical facility:

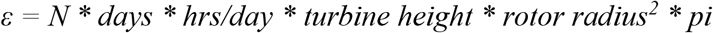

Where *N* is the number of turbines. We assumed turbines were 100 m high with 50 m blades and operated for 10 hr/day for 365 days. We evaluate changes in discrepancy as a function of inherent eagle activity rate, and survey effort using a constant facility size of 200 turbines.

The collision risk model guards against situations where surveys fail to detect eagles that are in fact present around the site, which means that sites where there are truly no eagles will have some predicted take. Therefore, we also evaluated estimate discrepancy as a function of survey effort at sites with no eagle activity across a range of facility sizes by varying the number of turbines from {50, 100,…, 500}.

### Cost analysis

We estimated the relationships between total eagle permitting costs and survey effort across a range of facility sizes and eagle activity rates. Total cost was calculated as the cost of preconstruction surveys for eagle activity and the cost of required mitigation based on predicted fatalities. Pre-construction survey cost estimates were provided by Western EcoSystems Technology, Inc. These ranged from $4,000 - $10,000 for the FWS minimum 24 hours of survey time at a single cylindrical plot over two-years. These rates include costs associated with the actual survey, travel to and from the facilities, data analysis, reporting and facility administration. In order to estimate a continuous relationship between cost and survey effort, we converted these to hourly rates per survey plot ($167/hour and $417/hour, respectively).

Compensatory mitigation costs (hereafter “mitigation”) are most commonly paid in the form of power pole retrofits that prevent eagle electrocution (U.S. Fish and Wildlife Service 2016). The cost per eagle varies depending on the lifespan of a retrofit and the presumed baseline eagle electrocution rate. We use estimates from a report estimating the per eagle cost and efficacy of retrofits (Hosterman & Lane 2017) for the Service. The low and high cost estimates to mitigate the take of one eagle through power pole retrofits were $15,200 and $38,000, based on a per-pole cost of $1,040 and $2,590, respectively, assuming an offset rate of 0.0051 eagles/year*pole and an active lifespan of 30 years for each retrofit (Hosterman & Lane 2017).

We used the same set of simulated scenarios and range of facility sizes to estimate the relationships between costs, survey effort, eagle activity rate, and facility size. For each combination of activity rate and facility size, we calculated survey, mitigation, and total costs across our simulated range of survey efforts and identified the amount of survey effort that produced the minimum total cost. This analysis was conducted for each combination of low and high mitigation and survey cost estimates, for a total of four cost optimization analyses.

## Results

### Model behavior

Discrepancies between eagle fatality estimates predicted with and without prior information increased at extreme high and low values of eagle activity (Fig. 1). For scenarios with eagle activity rates < 1.75 min/hr*km^3^, predicted fatalities estimated using priors were higher than site-specific estimates (i.e., above the 1:1 line in Fig. 1a). For scenarios with eagle activity rates > 1.75 min/hr*km^3^, predicted fatalities estimated using prior exposure information were less than site-specific estimates (i.e., below the 1:1 line in Fig. 1a). The more distant simulated eagle activity rates were from this inflection point, the greater the estimate discrepancy.

**Figure 1.**
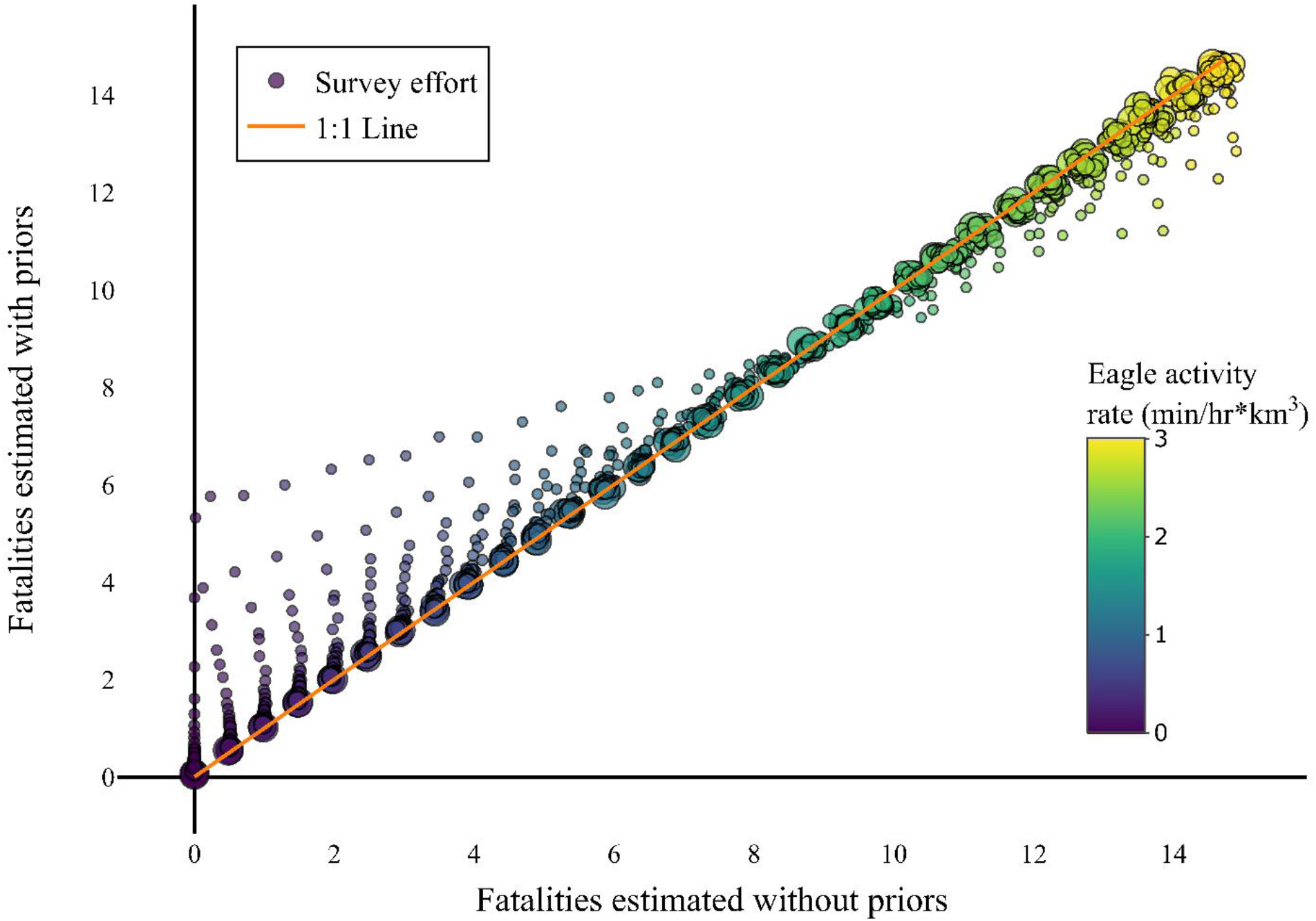
The collision risk model can both over and underpredicts eagle fatalities relative to survey data. Whether the model over or underpredicts fatalities is determined by whether eagle activity at a facility is above or below the mean of the prior eagle exposure distribution. Plotted values are the 80th percentile of posterior fatality distributions estimated for different levels of eagle activity and survey effort at a facility with 100 turbines 100 m high with 50 m rotor radius.

Greater survey effort reduced fatality estimate discrepancies (Fig. 2). The magnitude of the effect of effort on estimate discrepancy was contingent upon the observed eagle exposure rate (Fig 2a) and the size of a facility (Fig 2b). At the Service recommended minimum amount of survey effort (9.648 hr*km^3^), the absolute value of discrepancy was <1 eagle per year for all scenarios (Fig 1a). The slope of the relationship between survey effort and discrepancy (*Δ*) should be proportional to the difference between expected values of the posterior distributions calculated with and without priors.

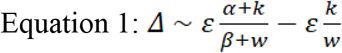

**Figure 2.**
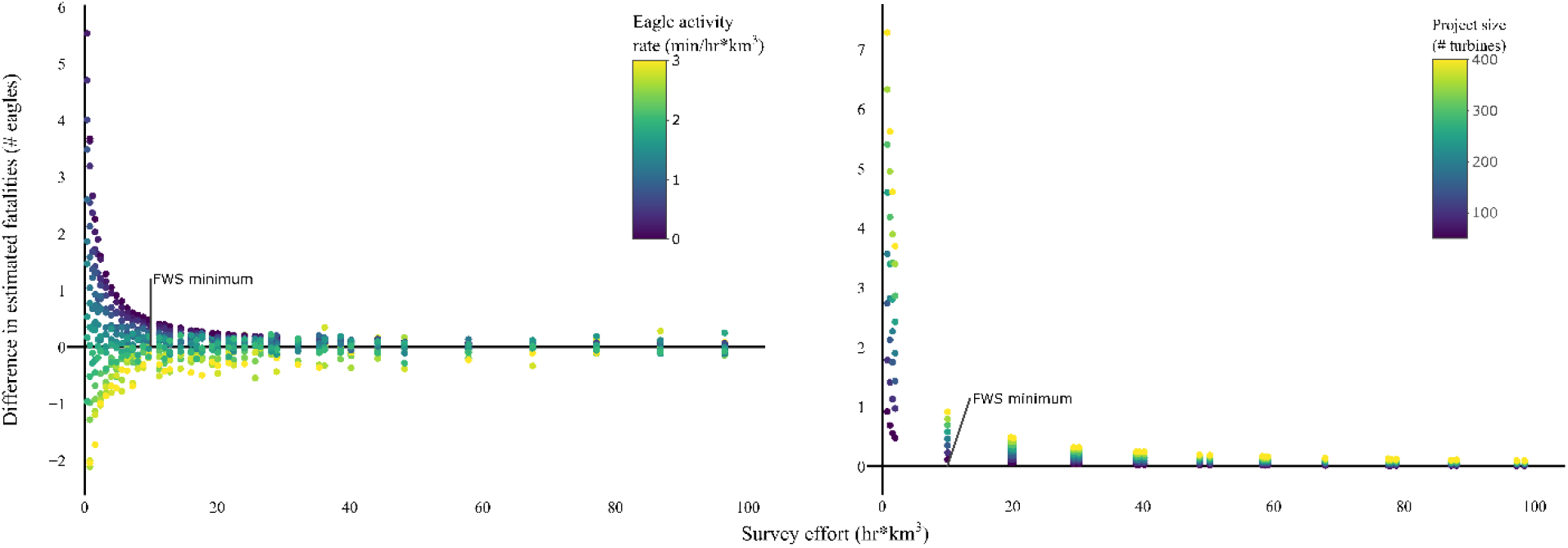
Survey effort decreases the discrepancy between eagle fatality estimates made with and without prior information. Plotted values are the difference in 80^th^ percentiles of the posterior distribution of eagle fatalities estimated with and without prior eagle exposure information given 100 m high turbines with 50 m rotor radius at a) facilities with 100 turbines and varying levels of eagle activity or b) facility with varying numbers of turbines and no eagles present.

Where *α* and *β* are the rate and shape parameters of the exposure prior, *k* is the observed number of eagles, *w* is survey effort and *ε* is the facility expansion factor. In our simulations the number of observed eagles, *k*, was equal to the product of eagle activity rate, *r*, and survey effort, *w*. We can substitute and simplify:

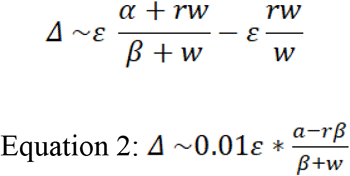

Equation 2 provides the expected discrepancy for a given exposure prior, survey effort, eagle activity rate, and facility specifications. Because the Service uses the 80th percentile, rather than the mean, of the posterior distribution this equation is not exact. However, it provides an empirically good model of these relationships, as indicated by *post hoc* measures of accuracy and bias between observed and predicted discrepancies within our simulated data (*RMSE* = 0.12, *MSD* = 0.012, *R^2^* = 0.98), although a Breusch-Pagan test revealed heteroskedasticity in the predictions (*BP1* = 119.27, *p* < 0.001).

We used Equation 2 to calculate the effort needed to effectively reduce the discrepancy in predicted fatalities to 0 over a five-year permit (i.e. Δ <= 0.1/year):

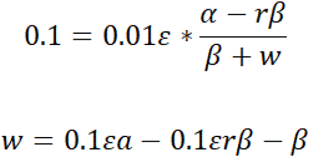

If the true eagle activity rate is zero, the effort needed to obtain zero predicted fatalities over five years is:

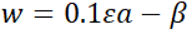

As seen from Equation 2, the degree of the response is dependent on the size of the facility. In scenarios with no eagle activity the size of the facility determined the effort required to obtain zero predicted eagle fatalities (Fig. 2b). The Service-recommended minimum survey effort reduced fatality estimates to 0.1 eagles/yr at the smallest facility considered (50 turbines) – effectively zero eagles for a five-year permit - and 0.9 eagles/yr at the largest facility considered (400 turbines). For the largest facility size 111 hr*km^3^ of survey effort, or 275 survey hours, was required to reduce predicted fatalities to zero over a five-year permit (i.e., < 0.1 eagle/year).

### Survey vs. mitigation costs

The relationships between survey effort and eagle take permitting costs varied predictably depending on eagle activity rate and facility size. For scenarios with eagle activity less than the prior exposure distribution mean, more survey effort reduced total costs to a point (Fig 3). Minimum cost occurred when the posterior estimate was effectively determined by site-specific survey data, after which costs increased linearly with additional survey effort (Fig. 3). In scenarios with high eagle activity (i.e. > 1.75 min/hr*km^3^) any amount of survey effort increased costs (Fig. 4). The amount of survey effort at which minimum costs occurred depended on the size of a facility. Smaller facilities achieved minimum total cost with less survey effort than larger facilities (Fig. 4). In general, the amount of survey effort that minimized costs increased with facility size and decreased with eagle activity rate (Fig. 4).

**Figure 3.**
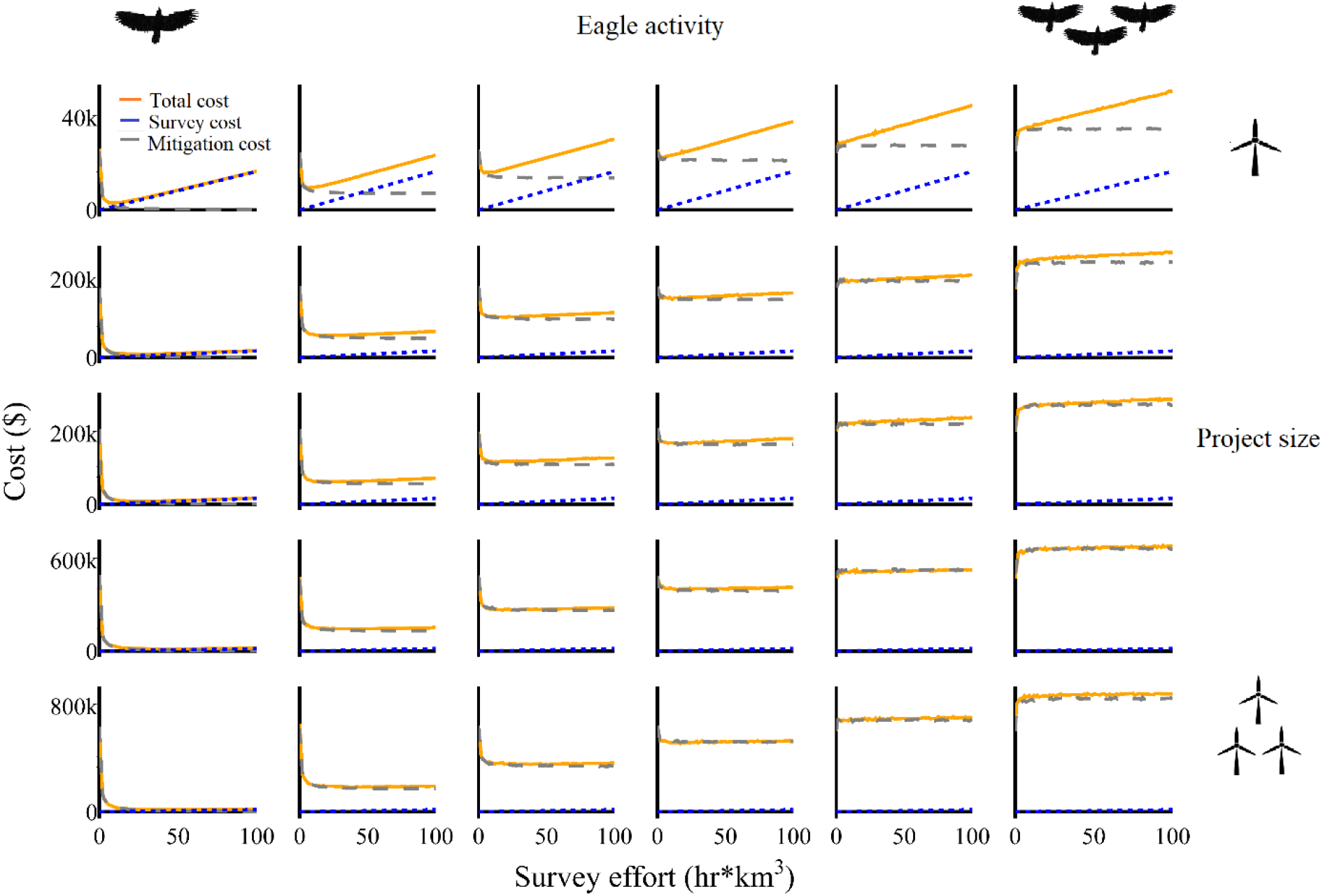
Additional survey effort can decrease costs associated with eagle take permitting. Curves show total cost (orange), which is comprised of survey costs (blue) and mitigation costs (grey) resulting from predicted eagle fatalities at different simulated combinations of eagle activity rate and facility size. Curves were generated using the low survey ($167/hr) and low mitigation ($1,040/retrofit) cost estimates.

**Figure 4.**
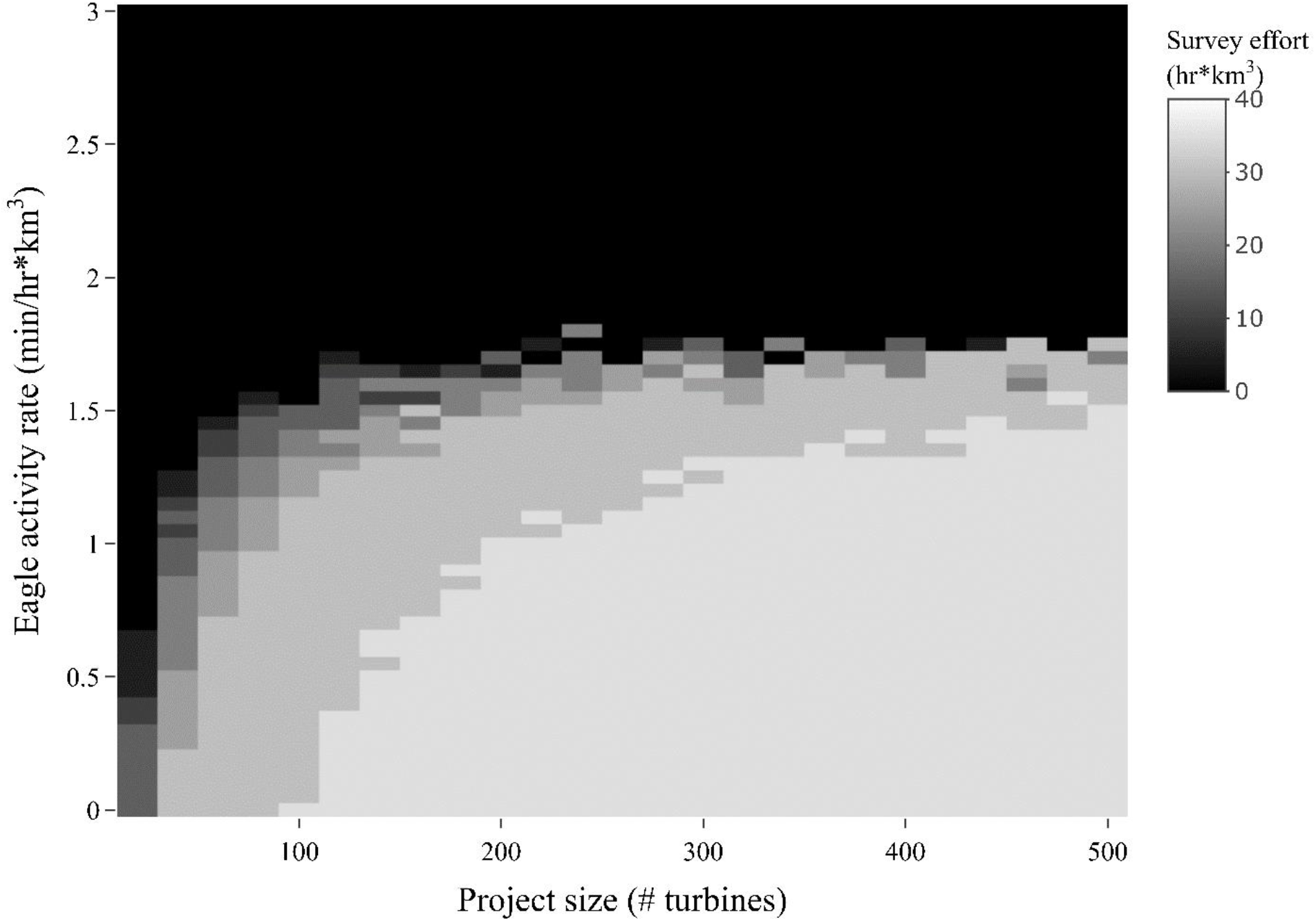
The level of survey effort resulting in minimum total cost (surveys + mitigation) varies as a function of facility size and eagle activity rates. Heatmap values represent the survey effort at which minimum total costs are achieved for a wind facility with given eagle activity rates and size. Turbines are assumed to be 100 m tall with 50 m rotor radius.

## Discussion

The development of renewable energy is an important component of biodiversity conservation, and policies are needed that align these two endeavors. This simulation study demonstrates and clarifies three important characteristics of the collision risk model used by the Service to estimate eagle fatalities at proposed wind energy facilities. First, the incorporation of prior eagle exposure information can both over- and under-estimate eagle fatalities relative to the inherent characteristics of a site. Each situation occurs predictably depending on whether true eagle activity at a site is above or below the mean of the prior exposure distribution (1.75 min/hr*km^3^). Second, the amount of time and area surveyed for eagle activity determines the strength of influence exerted by prior exposure distributions on final estimates. This behavior intuitively makes sense and is a desirable product of a Bayesian framework – at sites for which extensive information on local eagle activity exists, estimates of eagle fatality should more closely reflect that data. Finally, following from the interaction of these two findings, our analysis of survey and mitigation costs identified predictable relationships between survey effort, facility characteristics, and the costs associated with obtaining an eagle take permit. From these relationships we can predict the minimum costs for a range of known eagle activity rates and facility sizes – information that could be used to develop a policy or set of policies for issuing eagle take permits more expeditiously and with greater certainty in qualifying situations.

One of the appeals of Bayesian modeling is the ability to incorporate prior information and uncertainty into estimates to help offset bias induced by limited sampling or lack of direct experimentation (Harwood & Stokes 2003). In the context of eagle take permitting at wind facilities, our results confirm that prior information influences posterior eagle fatality estimates (Bay et al. 2016). In the context of Bayesian analysis, a lack of data can lead to less informative or biased priors. The current prior exposure distribution was built using data from nine wind facilities (New et al. 2015), and wind energy developers have stated that the facilities used to build the current prior exposure distribution represent a limited and biased sample (Speerschneider 2018). Historically, a lack of standardization among monitoring protocols at wind facilities has limited the ability to analyze greater amounts of data (Conkling et al. 2020), and the Service has created standard criteria for data to be included in the development of priors (U.S. Fish and Wildlife Service 2013). The collection of a larger prior dataset would address one of the primary criticisms of the collision risk model from the wind energy industry. An expanded dataset would enable the construction of subsets of priors that can address important sources of variation in eagle exposure, such as by region. To this end, the Service has completed an analysis to update the collision risk model priors using data from 80 facilities (New et al. 2018).

The inclusion of prior information also led to both under and overestimation of eagle fatalities (relative to using survey data alone). In our simulated scenarios these outcomes occurred predictably depending on whether the simulated eagle activity rate for a scenario was greater or less than the mean of the prior exposure distribution (1.75 eagle minutes/hr-km^3^). The degree of this discrepancy was proportional to the discrepancy between the inherent eagle activity rate for a given scenario and the prior exposure distribution mean. While the actual value of this threshold is subject to change as prior distributions are updated with new survey data, the patterns will be consistent. As an example, the mean of the proposed new prior exposure distribution *Γ*(0.287, 0.237) is 1.21 min/hr*km^3^ – less than the current distribution mean (New et al. 2018). Using this prior distribution, the collision risk model would overestimate eagle fatalities at fewer sites, and the magnitude of these discrepancies would be less while potentially underestimating fatalities in a greater number of scenarios.

Regardless of the priors used, the degree of discrepancy in eagle fatality estimates is also determined by the amount of survey effort. With additional survey effort, fatality estimates for a given scenario more closely reflected the simulated level of eagle activity observed rather than the prior exposure distribution. At the Service recommended minimum survey effort (9.648 hr*km^3^), the degree of fatality discrepancy was less than one eagle per year across a wide range of simulated site conditions and facility types. By increasing survey effort beyond this minimum, the influence of prior exposure information on fatality estimates was further reduced and the discrepancy between fatality estimates decreased. For even the largest facilities simulated, it was possible to reduce discrepancies to 0.1 eagle fatalities per year – effectively 0 eagles over a five-year permit.

While wildlife conservation agencies are concerned that more eagles may be killed at a project than expected, a concern for wind energy developers is that they be required to mitigate for more eagles than would likely be killed at their facility. The overprediction of fatalities can generate substantial costs, as power pole retrofits may be $6,800 - $103,000 per eagle - depending on retrofit lifespan and assumed electrocution rate (Hosterman & Lane 2017). These costs could be decreased by improvements in the cost effectiveness of existing mitigation measures (e.g. power pole retrofit lifespan), or the development of effective low-cost mitigation approaches. Regardless of the exact cost of mitigation per eagle, the amount of pre-construction survey effort determines the amount by which permitees may ‘overpay’ for mitigation if eagle activity at a site is in fact below the mean of the prior exposure distribution. In simulated scenarios with no eagles present, even the largest facilities considered would be able to reduce annual estimated fatalities to ≤ 0.1 eagles after 273 hr*km^3^ of survey effort. The rate of discrepancy reduction as a function of survey effort is predictable given site and facility characteristics, and we identify an equation describing this relationship (Equation 2). By explicitly incorporating the costs associated with survey effort, such a relationship may serve as a guide to identify which sites and under what conditions additional survey effort is beneficial from both a cost, feasibility, and eagle conservation perspective.

Our cost analyses identify circumstances in which wind energy facility operators can save money through additional surveying by decreasing predicted take estimates, and those in which eschewing surveys for up-front mitigation may benefit eagles and developers. In all situations, the greatest ratio of eagles saved per dollar spent was achieved by paying into mitigation without surveying. For facilities at which eagle activity is likely to be near or above the prior mean, robust pre-construction survey efforts are needed to ensure eagle fatalities are sufficiently compensated through mitigation. For small facilities that are likely below the prior exposure distribution mean reduced survey requirements in exchange for baseline amount of mitigation. In this scenario permittees would benefit from regulatory certainty and an expedited process, and more money would go towards eagle conservation than if additional resources were spent on surveys. At larger facilities with eagle activity rates likely below the mean, permit seekers could have the option to mitigate up front or conduct additional surveys. Mitigation is more expensive at large facilities due to larger expansion factors amplifying predicted fatality rates, and total costs are therefore determined to a greater degree by predicted fatality rates. In these scenarios, developers are likely able to reduce the total cost associated with obtaining an eagle take permit and can make their own decision as to whether additional survey effort represents a worthwhile investment or if they prefer to pay a fixed amount.

The exact thresholds identified here are subject to the assumed survey and mitigation costs, and a caveat to the results we present are that these estimates were provided by one company, respectively. In addition, survey cost estimates implicitly combined a component that scales with survey effort, as well as invariant administrative components. This means that survey costs likely do not increase as rapidly with increased survey effort as we estimated. Similarly, mitigation costs may be higher in practice. Either of these cases would expand the range of scenarios for which additional pre-construction surveys would reduce costs. Additionally, this approach would require some a priori indication of the eagle activity present at a site. One option would be to use national occupancy or abundance datasets like the Cornell occupancy model for bald eagles (Fink et al. 2010) or the Service’s spatially explicit golden eagle exposure models (Bedrosian et al. 2020) as a representation of eagle activity rate. These models are already being considered by the Service as a proximate measure of eagle activity as part of a proposed low risk eagle take permitting alternative. The alternative process would exempt developers proposing facilities in areas of low eagle activity – identified using percentile thresholds - from conducting pre-construction surveys (U.S. Fish and Wildlife Service 2019). Such an approach may also need to consider the spatial overlap between eagle use and wind resources. Regardless, our results could inform an alternative delineation of lower risk sites based on the characteristics of sites for which additional surveying reduces eagle fatality estimates relative to the prior exposure distribution mean.

Alternatively, the relationships quantified here could be used in a decision theory framework to generate policies that minimize costs for wind energy developers and maximize conservation benefits to eagles. Approaches such as Markov decision processes and value of information theory can be used to produce a recommended set of actions that maximize an objective while accounting for costs given different scenarios. Models like these have been used to determine optimal conservation policies balancing tradeoffs between monitoring and management action (Chadès et al. 2008; Bennett et al. 2018), and in situations with multiple variables and levels of uncertainty (Bal et al. 2018). As a prerequisite, these approaches require probabilities and costs associated with transitioning between ‘states’ of a system. In the case of eagle take permits we can use Bayesian posterior exposure distributions and survey and mitigation costs. Additionally, input from industry experts regarding the value of permit expediency could be used to realistically incorporate costs associated with delays due to additional surveys. Such an approach could provide a rigorous and defensible way for both conservationists and regulated entities to define thresholds for surveying and mitigation that optimally balance cost minimization and the maximization of conservation benefits to eagles.

This study effectively demonstrates the predictable patterns in the behavior of the collision risk model, and the application of simulation results to applied scenarios requires context. For simulated scenarios, we generated the number of eagles observed during hypothetical surveys deterministically based on a ‘true’ rate of eagle activity at the site. In situ, any given survey outcome will represent a random observation from an underlying activity distribution (e.g. Poisson) with a mean rate. We could have used the same set of simulated scenarios but drawn observed eagle exposure from a Poisson distribution defined by the simulated mean activity rate. We did not take this approach because our goal was to provide guidance on the general behavior of the model and introducing this stochasticity could have obfuscated those findings. Similarly, all simulations assumed a facility comprised of turbines with 100 m hub height and 50 m rotor radius, and absolute estimates of discrepancy and costs for a given facility are subject to different specifications. The relationships identified between survey effort, eagle activity rate, estimate discrepancies, and costs will remain. Finally, we focused our analysis on the direct financial costs of survey and mitigation efforts and acknowledge these activities can carry other costs to developers. Permitting delays construction due to the Service advocated minimum two-years spent surveying for eagles (U.S. Fish and Wildlife Service 2013) as well as other meetings. Unless wind energy developers also own power lines, they must find partners wiling to enact retrofits – often within the same geographic region as the proposed project – which can pose a barrier to implementing mitigation requirements. Developers likely take these costs into consideration when contemplating the benefits of pursuing a take permit.

A holistic evaluation of the impacts of renewable energy on wildlife must consider not only the direct contribution of renewables to wildlife mortality, but their contribution relative to other energy sources (e.g. coal, nuclear, natural gas). Renewable energy sources – particularly hydropower and wind – are relatively well studied regarding impacts on wildlife, in part because these effects are direct, observable, and highly publicized. Conversely, the mortality impacts of non-renewable sources have been less intensively studied, and have diffuse, indirect effects on wildlife (Loss et al. 2019). Additionally, the development and adoption of new technologies, such as automated real-time monitoring and curtailment (McClure et al. 2018) have the potential to further reduce mortalities associated with wind energy operation. Therefore, in addition to the benefits generated by curtailing climate change, the displacement of fossil fuel energy production with renewable sources likely brings other positive impacts for wildlife. While these benefits are difficult to model and incorporate into a permitting program, they should be considered when creating conservation policies pertaining to the development of renewables.

Ultimately, the eagle take permitting program under BGEPA is voluntary (U.S. Fish and Wildlife Service 2013). There is no requirement for wind energy facility operators to contribute to eagle conservation by obtaining eagle take permits - only legal consequences if they kill eagles without a permit. Thus, while an initial approach to eagle conservation might be to require wind facilities to take extensive measures to guarantee all eagle take is compensated for, an opaque or onerous process may incentivize developers to eschew permitting entirely. A permitting process that requires extensive effort to guarantee optimal outcomes may deliver lesser conservation benefits than one that accepts tolerable outcomes in the face of uncertainty (Burgman et al. 2005). By identifying requirements that balance regulatory certainty and expediency with reasonable protections, more can be done on the landscape to benefit eagles. The results of the analyses presented here can be used to identify permitting requirements that optimize a balance between eagle protection and renewable energy development.

